# Widespread genetic testing control inherited polycystic kidney disease in cats

**DOI:** 10.1101/2024.12.15.628535

**Authors:** Hisashi Ukawa, Akane Kida, Kai Ataka, Ryo Horie, Yuki Matsumoto

**Author notes:** **Corresponding author: Email:**; **Address:** 1-17-71 Fuchinobe, Chuo-ku, Sagamihara-shi, Kanagawa 252-5201, Japan.

## Abstract

A genetic variant of *PKD1* in cats, which causes polycystic kidney disease (PKD), is a target for direct-to-consumer genetic testing to assess PKD risk; however, its effect on genetic structure in cat populations remains unexplored. Therefore, in this study, we aimed to examine the changes in feline PKD and the *PKD1* variant across breeds and over time using a large dataset of 110,325 insured and 61,968 genetic-tested cats from 14 breeds. Results revealed that the *PKD1* variant frequency significantly decreased, with a reduction of 42.6% before 2019 and after 2022. Systematic genomic analysis revealed no differences in genetic structure or inbreeding levels. The effective population size of cats with the *PKD1* variant decreased between points. Overall, these findings highlight the potential of direct-to-consumer genetic testing in promoting more optimized breeding practices and enhancing feline welfare.

## Introduction

Polycystic kidney disease (PKD) is the most frequently inherited kidney disease found in humans (1, 2), cats (3), dogs, and livestock (4). PKD is either inherited as autosomal dominant polycystic kidney disease (ADPKD) or autosomal recessive types. In cats, ADPKD is one of the most commonly inherited diseases, and a nonsense single-nucleotide variant on chromosome E3 (g.42858112 C>A on the felCat9 assembly) was identified as a causative variant of ADPKD in 2004 (3) (OMIA:000807-9685 (5), the conventional *PKD1* variant). The conventional *PKD1* variant is considered to have nearly complete penetrance in cats, causing ADPKD in heterozygous cats and embryonic lethality in homozygous cats (3, 6). In addition, most cases of feline PKD are caused by the conventional variant (approximately 95% (7)). Because of the late onset of the disease [7 years on average (8)], affected cats are often bred before clinical signs are observed (3). Several studies have revealed that Persian and Persian-related cats have a high prevalence of the variant (9–11); this variant has also been identified in other cat breeds, such as the Scottish Fold (12). However, researchers in most studies analyzed only several dozens of samples and/or the results were biased as limited records from hospitals were used, primarily secondary care data with relatively high numbers of sick animals (12). More comprehensive analyses with large numbers of cats from various sources are required to establish epidemiological statistics for feline PKD to elucidate risk factors and contribute to advancing veterinary medicine. To this end, in the current study, we aimed to assess feline PKD onset and its association with the conventional *PKD1* variant using a large dataset of 110,325 insured and 61,968 genetically tested cats from 14 breeds.

## Results

### Evaluating PKD trends for insured cats

To characterize PKD onset in various cat breeds and its effect using the conventional *PKD1* variant in the cat population, combined analyses were performed using insurance data and genetic/genomic data for 110,325 insured cats of 14 breeds (Supplemental Table S1). First, we confirmed PKD onset according to breed and sex. We found that only 4 out of the 14 breeds tested had PKD: Scottish Fold, Munchkin, American Shorthair, and Persian (Fig. 1A and Supplemental Table S2). All four breeds have been reported to show PKD onset (11, 12), and we did not identify significant differences in the onset among the breed pairs (Fisher’s exact test: *P* > 0.05, Supplemental Table S3). Although there was a tendency for the incidence to be more than twice as high in females than in males, no significant difference was found (Fisher’s exact test: *P* > 0.05, Supplemental Table S4) as suggested by previous studies (10, 11, 13). To further characterize feline PKD, we evaluated cats that have been insured for over 10 years, and the rate of PKD according to age at first claiming was estimated (Fig. 1B). Out of 12,589 cats insured for over 10 years, 21 were diagnosed with PKD (0.167%, median age: 5 years). No age-related changes were observed between 0 and 10 years [logistic regression for age and onset: 0.026 (SE: 0.076), *P* = 0.73], indicating no association between age and PKD onset. Although a previous study reported that chronic kidney disease, a main clinical sign of PKD, occurs after the age of 3 (7), our data indicates the onset of PKD in younger cats (0–2 years old). To investigate PKD trends over the past 10 years, the proportion of PKD (within 3 years after birth) was analyzed. We observed no trend (Cochran-Armitage trend test: X-squared = 0.80, *P* = 0.37, Fig. 1C), indicating that the allele frequency of genetic variants associated with PKD may not have changed significantly over the past 10 years.

**Figure 1.**
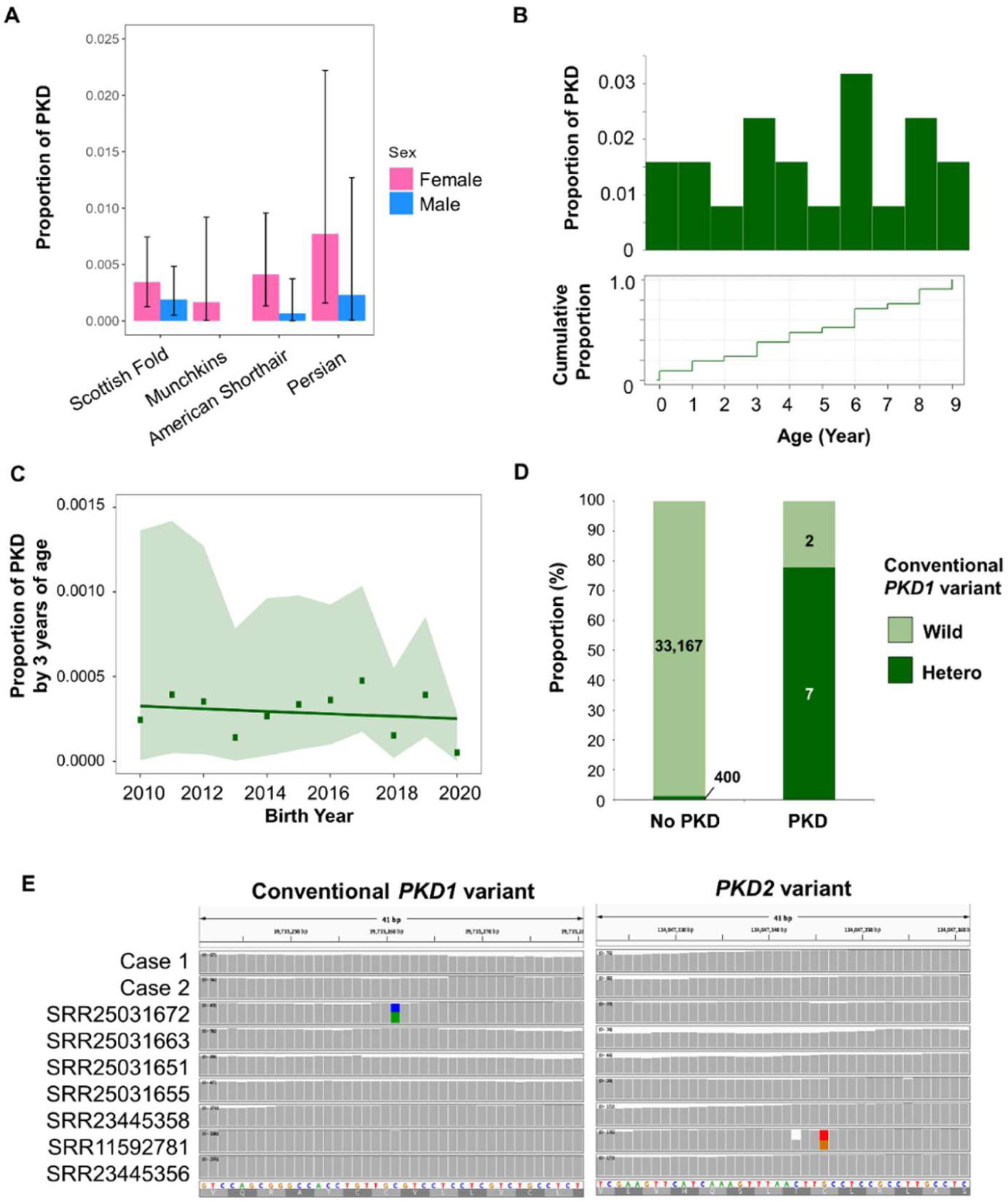
Characterization of feline PKD based on insured cat data. (A) Breed and sex differences in PKD onset using cats insured for over 10 years for four breeds that had a PKD claim. (B) Distribution of age of onset using cats insured for over 10 years for 4 breeds. Top: frequency based on histogram. Bottom: cumulative proportion by age. (C) Frequency of PKD found in cats insured for 3 years (14 breeds). (D) Comparison between conventional *PKD1* variant type and PKD onset in 14 breeds. (**E**) IGV view indicates no variant in cases 1 and 2 potentially associated with PKD, as revealed via whole-exome sequencing.

We evaluated the association between PKD claims and the *PKD1* variant among 33,576 cats who were insured during the last 3 years. We found a significant difference between the presence of the conventional *PKD1* variant and the occurrence of PKD (Fisher’s exact test; *P* < 1.29e-12, Fig. 1D). We found a 1.2% incidence of the conventional *PKD1* variant in non-PKD cats insured for 3 years, suggesting that these cats may develop PKD in the future. The analysis revealed that seven out of the nine PKD-affected cats had the conventional *PKD1* variant (77.8%) and that two did not have the variant. To uncover genetic variants associated with PKD other than the conventional *PKD1* variant, exome sequencing data for the two cats (cases 1 and 2) and additional open data of whole genome and exome sequencing were analyzed (n = 104 including seven PKD-affected cats, Supplemental Table S5). Upon searching 92 out of 121 genes associated with human PKD, which are orthologs between human and cat genomes, we identified 10 variants, including the conventional *PKD1* variant, as candidates causing PKD (Supplemental Table S6). Three potential variants in *PKD1*, in addition to the conventional *PKD1* variant, were detected in PKD-affected cats in the database. In addition, we identified six high-impact variants in the *IFT80*, *ANKS6*, *RPGRIP1L,* and *PKD2* genes, and a variant in the *PKD2* gene, which is a frameshift variant that was recently detected (14). However, no variant was detected in both cases from insured cats (Fig. 1E), suggesting that other unknown genetic factors such as cis-regulatory elements and/or epigenetic factors might be associated with PKD onset (7). Several studies have also reported cats without the conventional *PKD1* variant, revealing PKD-positive histological evidence and indicating the possibility of other variants causing similar pathologies (7, 12, 15). Therefore, further research, including more comprehensive datasets obtained by pan-genome approaches, is required to genetically and epigenetically characterize feline PKD.

### PKD1 variant trajectories across breeds and years

The allele frequency of the conventional *PKD1* variant has been investigated worldwide, including the USA (3), Mexico (16), UK (15), Switzerland (17), Slovenia (9), Taiwan (13), Turkey (18), and Japan (11, 12). Most studies conducted thus far have been limited to cats that visited animal hospitals; therefore, the prevalence of the conventional *PKD1* variant may be biased. Samples for genetic studies should be collected from diverse populations and breeds (19). In addition, direct-to-consumer (DTC) genetic testing of cats has been recently introduced; however, its effects on the allele frequency of risk variants remain unclear.

To clarify the frequency of the conventional *PKD1* variant in cats, we used a large dataset of 61,968 cats from 14 breeds as well as the effect of widespread DTC genetic testing on the variant prevalence and inbreeding over time. Real-time PCR was performed to determine the cat genotypes. In 2022, no cats harboring the conventional *PKD1* variant were found among Ragdoll, Main Coon, Bengal, and Russian Blue; however, the *PKD1* variant was found in the eight remaining breeds (Fig. 2A and Supplemental Table S7). For the eight breeds, the conventional *PKD1* variant frequency ranged between 0.99% (British Shorthair) and 14.07% (Himalayan) in 2019 and between 0.71% (American Shorthair) and 9.30% (Himalayan) in 2022. The overall frequency of the *PKD1* variant in our study was 2.08% among the 62,304 cats born in 2019 or 2022. We identified no cats with homozygous *PKD1* variants in the datasets, suggesting that this variant is embryonic lethal (3, 7).

**Figure 2.**
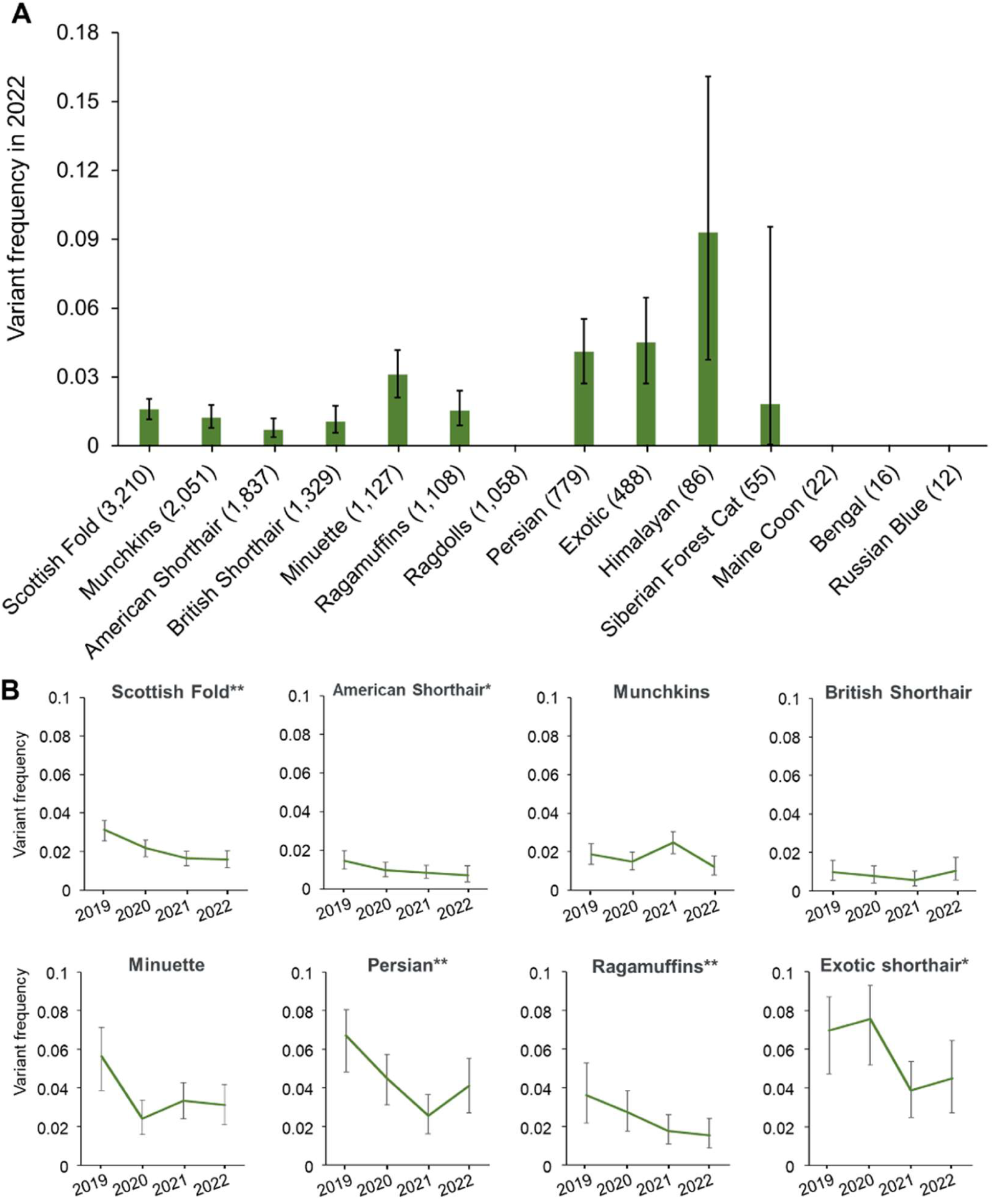
Frequency of the *PKD1* variant. (A) Frequency of the *PKD1* variant in 22 cat breeds born in 2022. The number in parentheses indicates the number of genotyped cats. (B) Trajectory of allele frequency of the *PKD1* variant in eight breeds with larger samples in each year (n > 100). * and ** adjacent to breed names indicate suggestive and significant decreases, detected via a binomial test.

Among the eight breeds analyzed using the logistic regression model, the proportion of the conventional *PKD1* variant decreased in three breeds of cats born in 2019 or 2022 (Scottish Fold, Persian, and Ragamuffins; significant differences) and two breeds (American Shorthair and Exotic Shorthair; suggestive differences) among the eight breeds (Fig. 2B and Supplemental Table S8). No significant or suggestive changes were observed in the remaining three breeds (Munchkins, British Shorthair, and Minuette). These results indicate that genetic testing promotes a decrease in conventional *PKD1* variants, although its effectiveness might vary with the breed. We identified no critical factor that explains the observed differences; however, the intent of cat breeders and/or diffusion of genetic testing depending on the cat breed are potential factors. The conventional *PKD1* variant should be eradicated via careful breeding with genetic monitoring to avoid the spread of the variant in all breeds with the existence of the variant, especially for breeds showing no changes, such as Minuette, and high-prevalence breeds, such as Himalayan.

### Effect of genetic testing on the cat genome

Genome-wide single-nucleotide polymorphisms (SNPs) were assessed in 81 cats to clarify the genetic structure (Supplemental Table S9), genetic diversity, and effective population size at two time points: before genetic testing started (2019) and after genetic testing became popular (2022). We analyzed the genetic structure through clustering using ADMIXTURE and principal component analysis (PCA) for two breeds: Scottish Fold and Persian. These two breeds were selected because of the significant decrease in the *PKD1* variant following the spread of genetic testing and because relatively large sample sizes could be included to reduce sampling bias. The clustering results showed that the cross-validation error was the lowest when *K* = 1 in both breeds, indicating no structure in the compared populations. Similarly, PCA also identified no obvious components that explained the variation in the cat group or birth year (Fig. 3A, B). Furthermore, no clear clusters were identified in the neighbor-joining tree (Fig. 3C, D). We compared the inbreeding coefficients between the groups based on runs of homozygosity (*F*_ROH_; Fig. 3E, F). The mean *F*_ROH_ of the Scottish Fold breed was 0.10 ± 0.081 (SD), 0.10 ± 0.078, 0.06 ± 0.046, and 0.06 ± 0.026 for the wild-type cats born in 2019 (Wild 2019), cats with the conventional *PKD1* variant born in 2019 (Hetero 2019), wild-type cats (Wild 2022), and cats with the conventional *PKD1* variant born in 2022 (Hetero 2022), respectively. The mean *F*_ROH_ of the Persian breed was 0.11 ± 0.029 (SD), 0.12 ± 0.045, 0.11 ± 0.038, and 0.11 ± 0.034 for the Wild 2019, Hetero 2019, Wild 2022, and Hetero 2022 groups, respectively. Overall, no significant differences were detected (Fig. 3A, B and Supplemental Table S10). In addition, the observed heterozygosity of the two breeds was computed for the four groups to investigate genetic diversity. We discovered no differences among the four groups of the Persian breed (Supplemental Table S11).

**Figure. 3.**
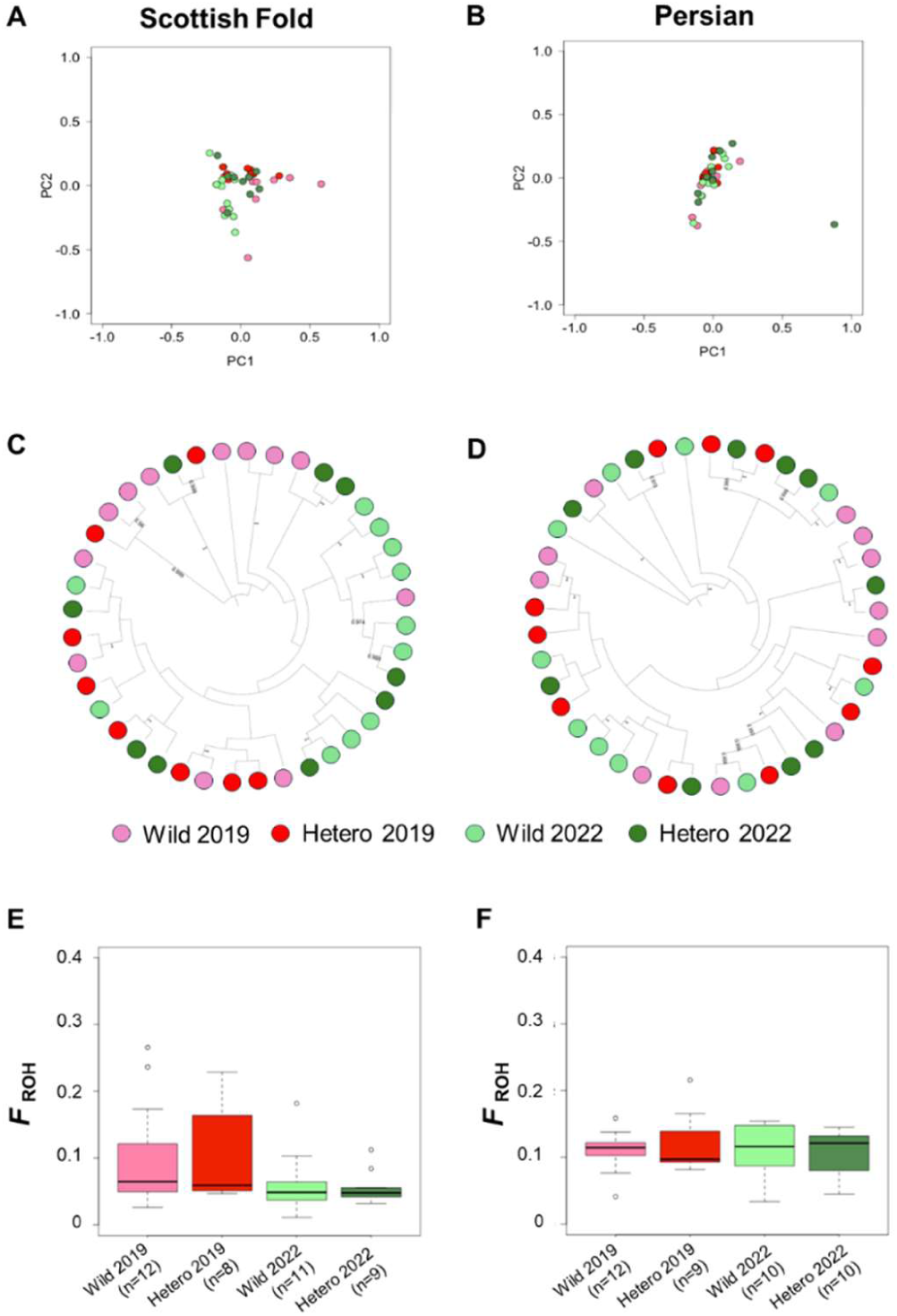
Genetic relationships and inbreeding levels of two breeds. (**A**, **B**) Principal component analysis (a: Scottish Fold, b: Persian). (**C**, **D**) Phylogenetic tree based on the neighbor-joining method (c: Scottish Fold, d: Persian). The percentage of replicate trees in which the associated taxa clustered together in the bootstrap test (1,000 replicates) is shown next to the branches. Only over 95% replicates are shown. Inbreeding coefficient based on runs of homozygosity (**E**: Scottish Fold; **F**: Persian). No significant difference was observed in any of the comparisons.

To estimate the potential number of cats bred, we used the linkage disequilibrium method to determine the contemporary effective population size, *Ne* (Fig. 4). In the Scottish Fold breed, we computed the contemporary *Ne* of the Wild 2019 group at 375.3 (95% CI: 367.3–383.7), which was lower than that of the Wild 2022 group at 563.7 (95% CI: 546–582.4), whereas the *Ne* of the Hetero 2019 group was 475 (95% CI: 462.1–488.6), which was higher than that of the Hetero 2022 group at 202.5 (95% CI: 200.1–204.9). These results indicate that the number of cats bred with the *PKD1* variant decreased after genetic testing had been widely used in the Scottish Fold population. In contrast, in the Persian breed, the *Ne* of the Wild 2019 group was 119.2 (95% CI: 118.4–120), which was higher than that of the Wild 2022 group at 76.4 (95% CI: 76.1–76.7), whereas the *Ne* of the Hetero 2019 group was 208.7 (95% CI: 206.3–211.1), which was higher than that of the Hetero 2022 group at 70.1 (95% CI: 69.8–70.4). Estimation of the effective population size suggested that the decline in *Ne* was greater in the population with the conventional *PKD1* variant. Collectively, systematic genomic analyses suggested that breeders are reducing the number of cats carrying the conventional *PKD1* variant in these two breeds by avoiding inbreeding.

**Figure. 4.**
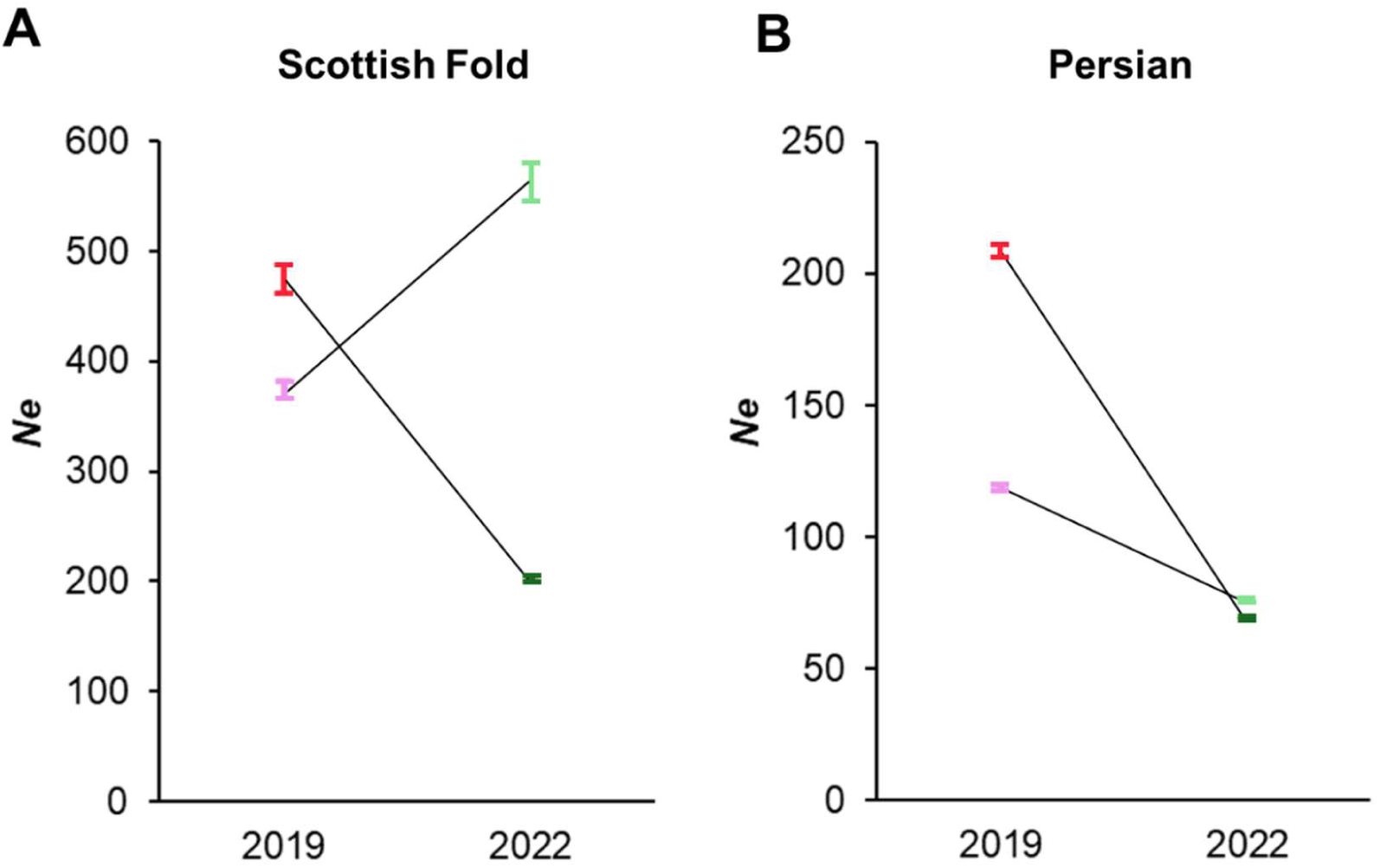
Effective population size (*Ne*) estimated using the genome-wide SNP data. Pink, red, light green, and dark green indicate Wild 2019, Hetero 2019, Wild 2022, and Hetero 2022, respectively. Error bar indicates 95% CIs for *Ne*. CI, confidence interval.

## Discussion

Genetic testing for disease-causing mutations in companion animals is increasingly being performed by veterinarians for diagnosis, by breeders for reducing the incidence of inherited diseases, and by pet owners for determining the genetic background of their pets (20). Therefore, accurate epidemiological information on the conventional *PKD1* variant frequency is useful for conducting genetic testing. The prevalence of the *PKD1* variant has been investigated over the past few decades, with the Persian breed being associated with a relatively high prevalence in Japan [46% (11) and 10.81% (12)]. In contrast, our results indicate a low prevalence of the variant in the Persian breed (4.11–6.72%). Previous studies used cats that visited university veterinary teaching hospitals as the sampling pool; therefore, the collection was likely to be biased toward unhealthy cats. However, in the current study, we used a more general population for data collection, which likely contained less biased epidemiological data.

The prevalence of the conventional *PKD1* variant varies among breeds as well as birth years, as demonstrated in our study. Population genetic studies of cats have shown that the genetic structure varies among countries (21); therefore, the *PKD1* variant frequency could vary among countries, in addition to breeds or birth years. Our results suggest that no Ragdoll cats harbored the *PKD1* variant among the 3,101 cats tested. Furthermore, the *PKD1* variant was not found in Ragdoll cats in Japan (12) and Belgium (22). To date, no *PKD1* variant has been confirmed in Ragdoll cats outside of the USA (3, 23). In contrast, we observed a higher variant frequency in the Himalayan breed in Japan. In Mexico and Taiwan, the *PKD1* variant has not been identified in any of the Himalayan cats; however, only a small number of cats have been tested (13, 16).

Moreover, previous studies have identified *PKD1* variants in randomly bred cats, in addition to inbred cats (11, 12, 15). Considering the existence of this variant in diverse cat breeds and randomly bred cats in several countries, the origin of the *PKD1* variant remains unknown. Therefore, future studies are necessary to identify the genetic origin of this variant to reveal the differences in prevalence across breeds and countries.

Using systematic genomic analysis, the present study highlights the value of genetic testing as a tool to lower the risk of ADPKD while avoiding inbreeding, as previously reported in a *SOD1* gene variant associated with canine degenerative myelopathy (24). The onset of PKD occurs at approximately 5 (this study) or 7 years of age (8). A relatively small percentage (approximately 5% (7) to 22.2% in this study) of cats develop PKD in the absence of the *PKD1* variant; therefore, most cases of ADPKD in cats are caused by the *PKD1* variant. A limitation of our study is that we could not detect a decrease in the prevalence of PKD. Considering the start of widespread DTC genetic testing in Japan, the corresponding decrease in the prevalence of the *PKD1* variant should be noticeable in the late 2020s. Most inbred cats are bred by humans, and some genetic variants, including the conventional *PKD1* variant, are considered to have nearly complete penetrance in cats (3, 6); therefore, to improve animal welfare, breeders should control the risk of such inherited diseases using genetic testing and desired breeding practices based on the results (11, 25).

## Methods

### Ethics statement

All swab samples were obtained with the consent of the owners. This study was approved by the Ethics Committee of Anicom Specialty Medical Institute, *Inc.* (*ID*: 2020-06, 2022-02, and 2024-01).

### Insurance data analysis

We used insured cat data to characterize PKD in a cat population, based on three perspectives: breed difference, sex difference, and age difference. Biometric information and insurance claim data were collected for cats that have been continuously insured to the pet insurance by Anicom Insurance Co., one of the largest pet insurance companies in Japan, from April 1, 2008 to December 31, 2023. We extracted and analyzed data of 12,589 cats for a period of more than 10 years (Supplemental Table S1). Individuals diagnosed with PKD or renal cysts and for which insurance claims were filed were classified as the PKD onset group, and individuals for which insurance claims were filed for other diseases, including renal diseases other than polycystic kidney disease, were classified as the non-onset group. The age at onset was the age on the day that PKD was registered in the insurance claim data. The incidence rate by onset age was calculated by dividing the number of individuals who developed the disease at age n by the total number of individuals minus the number of individuals who developed the disease by age n-1. A cumulative density function was calculated for the incidence rate by the onset age obtained. The differences among breeds and sex were analyzed using Fisher’s exact test with the *fisher.test* function in R version 4.3.3. We used logistic regression analysis with the *glm* function in R to investigate changes in the proportion of PKD by age.

To investigate the association between birth year and PKD proportions, we used 110,325 insured cats within 3 years of age. The number of PKD cats was counted by birth years and divided by the total number of cat births in the year. The trend of the proportional change by birth year was assessed using the Cochran-Armitage trend test using the *prop.trend.test* function in R.

To investigate the association between the variant type of the conventional *PKD1* variant (see methods details below) and PKD onset, we used 33,576 insured cats with genetic testing between January 1, 2017, and December 31, 2023. The number of PKD cats with variant data for the conventional *PKD1* variant was counted by the above criteria. The difference between PKD onset and variant type was analyzed using Fisher’s exact test using the *fisher.test* function in R.

### Next-generation sequencing data analyses

Two PKD-affected cats (cases 1 and 2) with no conventional *PKD1* variant and 104 cats from the sequence read archive were targeted with a genome-wide survey to uncover the potential variant associated with PKD (Supplemental Table S5). DNA samples from swabs of cases 1 and 2 were obtained. We used the Twist custom exome panel based on high-quality cat genome assembly (AnAms1.0) (26). Sequence libraries were generated using the manufacturer’s protocol and sequenced using the NovaSeq X Plus platform (Illumina, San Diego, CA, USA).

After fastq generation, the sequenced reads were mapped to AnAms1.0 using Illumina’s DRAGEN v 4.0.1. (Illumina), and joint calling using 106 cats was performed using the DRAGEN software. After generating joint-vcf, we annotated genes and the effect of the variant using SnpEff v5.1d (27). We referred to Kidney Cystic and Ciliopathy Disorders Gene Curation Expert Panel data sets in the ClinGen database, which is a National Institutes of Health-funded resource dedicated to building a central resource (https://clinicalgenome.org/affiliation/40066). First, 121 registered genes in the database were converted to AnAms1.0 gene sets based on its annotation, then 98 genes were detected. Additionally, all variants found in the 98 genes were filtered by three criteria: (1) impact HIGH by SnpEff annotation, (2) minor allele frequency < 0.01, and (3) only found in PKD-affected cats.

### Animals for genetic analyses

We obtained buccal swabs from 61,968 cats from 14 breeds (Scottish Fold, American Shorthair, Munchkins, British Shorthair, Minuette, Persian, Ragamuffins, Ragdolls, Exotic Shorthair, Himalayan, Selkirk Rex Longhair, Kincalaw, British Longhair, Siberian Forest Cat, Main Coon, Norwegian Forest Cat, Bengali, Somali, Russian Blue, Sphynx, Singapore, and Burmilla). All cats born between January 1, 2019, and December 31, 2022, were sampled by breeders, owners, and pet stores from around Japan. The breed and birth date data were collected from the owners.

### Genotyping of the PKD1 variant

DNA was extracted from oral mucosal tissue using commercial kits (chemagic™ DNA Buccal Swab Kit, PerkinElmer, Waltham, MA, USA; DNAdvance Kit, Beckman Coulter, Brea, CA, USA). Real-time PCR was performed to determine the genotypes of PKD-associated mutations (*PKD1* variant: g.42858112 C > A), specifically wild-type homozygotes (C/C, Wild), heterozygous carriers (C/A, Hetero), and variant homozygotes (A/A, Mutant), as previously reported (3). A real-time PCR genotyping assay was performed using TaqMan probes with fluorescein amidite (FAM) and 2’-chloro-7’-phenyl-1,4-dichloro-6-carboxyfluorescein (VIC) at one end and a non-fluorescent quencher (NFQ) at the other. The assay was performed using 5.0 μL of FastStart Universal Probe Master (Rox; Roche, Basel, Switzerland), 0.2 μL of Custom TaqMan SNP Genotyping Assay (Thermo Fisher Scientific, Waltham, MA, USA), >20 ng of DNA, and ultrapure water up to 10 μL. The Custom TaqMan SNP Genotyping Assay included forward (5ʹ-CCTCGGAGCCGCTTCAC-3ʹ) and reverse (5ʹ-TTGGCGCCCAGGAAGAG-3ʹ) primers and two probes, VIC-ACGAGGAGGACGCAACA-NFQ and FAM-CGAGGAGGACTCAACA-NFQ, for the wild-type and mutant alleles, respectively. Real-time PCR was performed on the fast cycle on a QuantStudio7FlexFast96Well Real-time PCR system (Thermo Fisher Scientific) for one cycle at 25 °C for 30 s and 95 °C for 10 min. This was followed by 40 cycles at 95 °C for 15 s, 60 °C for 1 min, and finally one cycle at 25 °C for 30 s. To determine the specificity of this assay, two samples each of wild-type and heterozygous mutant samples were selected randomly, and a water control was run on the genotyping assay. The genotype of each sample was determined by examining the amplification of the reporter dye (FAM or VIC) in an amplification plot.

To investigate the allele frequency trajectory of the *PKD1* variant over 4 years, we used eight breeds with a large sample size (i.e., over 100 cats/breed/year). A logistic regression based on the *glm* function in R was used, with the birth year of each breed as an explanatory variable and the numbers of wild-type and heterozygous individuals as response variables. The response variable was assumed to follow a binomial distribution, and the link function was a logit. The 95% CI for each genotype was calculated using the *binorm.test* function in R.

### SNP array data analyses

We used an SNP genotyping array for cats, the Feline 63 K SNP genotyping array (28) (Illumina), to investigate the effects of genetic testing on the entire genome. We compared genome-wide SNPs from four subpopulations of wild-type and heterozygous *PKD1* risk alleles, all derived from the Scottish Fold and Persian cats born in 2019 and 2022. These two breeds were selected because of the substantial decrease in frequency of the *PKD1* variant during the spread of genetic testing and because sufficiently large sample sizes could be obtained from a broad client base, thereby minimizing sampling bias. Considering the effect of genetic background on array-based analyses, we assessed all cats from the largest clients. We used an SNP genotyping array (Illumina Infinium iSelect 63K Cat DNA Array) and an Illumina iScan system to detect 63 K SNP genotypes. Genotype coordinates correspond to the genome assembly, Felis_catus_9.0.

Quality control was performed using PLINK version 1.90 (29). Missing rates for each locus (geno option) and individual cats (mind option) were both <5%. We did not examine the Hardy– Weinberg equilibrium, as we did not assume random mating for pedigreed cats. Population genetic analyses should not include close relatives (30); therefore, we conducted robust relationship-based pruning using the king-cutoff option in PLINK version 2.00a2LM (threshold of 0.176) to exclude pairings between monozygotic twins and first-degree relatives. For allele frequency-based quality control, we excluded SNPs with a minor allele frequency <0.01. We also removed sex chromosome variants and autosomal insertions-deletions because their effects on allele frequency differed from those of autosomal SNPs. The final analysis included 46,821 autosomal SNPs.

After quality control for individual cats, including removal of related animals, 81 cats remained: homozygous wild-type (C/C) cats tested in 2019 (Wild 2019, n = 24) and 2022 (Wild 2022, n = 21), as well as heterozygous cats tested in 2019 (Hetero 2019, n = 17) and 2022 from two breeds (Hetero 2022, n = 19; Supplemental Table S9). The subsequent analyses were performed using this dataset.

### Genetic structure analyses

To clarify the genetic structure of the cat population, we performed a maximum-likelihood ancestry analysis using ADMIXTURE version 1.3 (31) and PCA using PLINK version 1.9 with the default settings. We set the number of populations (K) between 1 and 10 for ADMIXTURE. We used option cv for the cross-validation error calculation to select the optimal *K* value based on the lowest error.

To estimate the genetic relationships within a breed, the neighbor-joining method was implemented in Molecular Evolutionary Genetic Analysis (MEGA) v 10 (32). The PLINK ped/map format described above was converted to the FASTA format using an in-house script. FASTA files were used as inputs to MEGA. The maximum composite likelihood method was used to compute the evolutionary distance. Tree construction was performed using the neighbor-joining method with 1,000 replicates using the bootstrap test.

### Estimating inbreeding coefficients and heterogeneity

We inferred the inbreeding levels using specimens obtained from genome-wide SNP genotyping. The R package DetectRuns was used to obtain the proportion of times each SNP fell within a run per population, corresponding to locus homozygosity or heterozygosity in the respective population. The following DetectRuns parameters were used: minSNP = 41, maxGap = 106, minLengthBps = 50000, and minDensity = 1/5000. Statistical tests were performed using the Wilcoxon test in the R software. Statistical significance was set at 0.05. We estimated observed heterozygosity for four groups (two genotype groups from two time points, 2019 and 2022) from two breeds (Scottish Fold and Persian) and their 95% CI using GenoDive v3.0.6 with default settings (33).

### Estimating effective population size

The contemporary *Ne* for each breed was estimated from the genome-wide SNP data using the linkage disequilibrium method in NeEstimator version 2.1 (34). We performed this analysis for four groups (Wild from 2019 and 2022 and Hetero from 2019 and 2022) of the two breeds, Persian and Scottish Fold.

## Data access

Exome sequencing data were deposited in DDBJ Bioproject (PRJDB19172). All SNP data have been deposited in the archives of the DRYAD database (https://doi.org/10.5061/dryad.6hdr7sr82).

## Acknowledgments

We acknowledge funding by the Research Foundation of Anicom Insurance Inc. (Japan), Anicom Specialty Medical Institute Inc. (Japan), and Anicom Pafe Inc. (Japan). We thank all the staff members performing genetic testing at Anicom Specialty Medical Institute Inc. and Anicom Pafe Inc. for their support. We would also like to thank Editage (www.editage.com) for English language editing.

## Author Contributions

Conceptualization: YM. Methodology: YM, HU. Investigation: YM, HU, AK. Visualization: YM, HU. Funding acquisition: YM, HU, KA. Project administration: YM, HU. Supervision: YM, HU, KA, RH. Writing – original draft: YM, HU. Writing – review & editing: YM, HU, AK, KA, RH.

